# Cholesterol depletion activates trafficking-coupled sphingolipid synthesis

**DOI:** 10.1101/2025.02.12.637879

**Authors:** Yeongho Kim, Jan Parolek, Christopher G. Burd

**Affiliations:** Department of Cell Biology Yale School of Medicine, New Haven, Connecticut 06520

**Keywords:** sphingolipid, cholesterol, coatomer II, lipid homeostasis, ceramide synthase, endoplasmic reticulum, Golgi apparatus

## Abstract

Within cellular membranes, sphingomyelin is associated with cholesterol and this complex facilitates homeostatic regulation of membrane viscosity. Acute cholesterol depletion increases the synthesis of very-long-chain (VLC) sphingomyelin, but a link between lipid sensing and sphingolipid synthesis is lacking. Using sphingolipid metabolic flux analysis, we observed that VLC-ceramide, the precursor to VLC complex sphingolipids that are produced in the Golgi apparatus, was rapidly consumed after cholesterol depletion, while synthesis of long-chain sphingolipids was unaffected. Sphingolipid trafficking assays showed that cholesterol depletion enhances VLC-Ceramide trafficking from the endoplasmic reticulum to the Golgi apparatus. Changes in the sizes of coatomer II ER exit sites were correlated with increased VLC-Ceramide trafficking and concomitant increase in sphingomyelin. Depletion of Sec16A, a component of the COPII network, abolished VLC-SM synthesis. This study reveals ER-to-Golgi trafficking of VLC-Ceramide as a key regulatory node in organelle membrane homeostasis pathways.

**Summary:** In cellular membranes, sphingomyelin is associated with cholesterol. Metabolic flux analysis of sphingolipid metabolism showed that synthesis rate of sphingomyelin, but not ceramide, was increased after depletion of cholesterol due increased rate of COPII-dependent ER-to-Golgi transport of ceramide.

## Introduction

The plasma membrane (PM) of the cell contains up to 80% of total cell cholesterol and sphingolipids (Lange et al., 1989). These lipids bind each other in organelle membranes and these complexes facilitate dense lipid packing and increased membrane tension. How cells maintain homeostatic levels of sphingolipids in the PM, and other organelles constitute a major gap in knowledge of organelle membrane homeostasis pathways.

Cholesterol is synthesized in the endoplasmic reticulum (ER), but the enzymes of complex sphingolipid synthesis pathways are distributed between the ER and Golgi apparatus, so these pathways require ER-to-Golgi trafficking of sphingolipid metabolites. Single lipid synthesis is initiated in the ER by synthesis of the long-chain sphingoid base (*e.g.*, sphinganine), which is the core of all complex sphingolipids. It is produced in the ER by serine-palmitoyltransferases (SPTLC), whose activity is regulated protein phosphorylation of negative regulators called ORMDL (Breslow et al., 2010). Subsequently, the sphingoid base is conjugated with a fatty acid by one of six ceramide synthases (CerS1-6) that differ in specificities for different fatty acid chain lengths (Cingolani et al., 2016). Ceramides are transported to the Golgi apparatus, where they are converted into complex sphingolipids, such as glycosphingolipids (GSL) and sphingomyelin (SM), through the addition of head groups in the *trans* cisterna and *trans*-Golgi network (TGN) of the Golgi apparatus (Pothukuchi et al., 2021).

At least two trafficking pathways transport ceramides from the ER to the Golgi apparatus: bulk membrane transfer and transport via the soluble ceramide transfer protein (CERT) that functions at the ER-Golgi membrane contact site (MCS). Bulk membrane trafficking of ceramide requires coatomer II (COPII), which is best characterized for its essential role in protein export from the ER, and we have shown that a sub-class of sphingolipids containing N-linked fatty acyl chains composed of more than 20 carbon atoms, termed very long chain (VLC) sphingolipids, accumulate in the ER when COPII function is reduced (Kim et al., 2023). CERT-dependent transport of long-chain (LC) ceramides (C≤20) continues unimpeded when the COPII pathway is blocked (Kim et al., 2023).

Metabolic regulation of sphingolipid levels is required to maintain plasma membrane lipid homeostasis in response to changes in membrane lipids (Kim and Burd, 2023). A study (Capasso et al., 2017) described a phosphatidylinositol 4-phosphate (PI4P) signaling network at the Golgi apparatus that constitutes a molecular rheostat that coordinately regulates changes in cholesterol and CERT-dependent LC-ceramide trafficking between the ER and Golgi apparatus. CERT and other LTPs are targeted to the ER-Golgi MCS, in part through recognition of PI4P (Hanada et al., 2003; Levine and Munro, 2002) that is produced on *trans*/TGN membranes (Balla et al., 2005; Wang et al., 2003). Much of the cholesterol in a cell is exported from the ER and transferred to the *trans*/TGN by oxysterol binding protein (OSBP), which exchanges bound cholesterol from the ER for PI4P from the Golgi membranes, lowering PI4P amounts on the *trans*/TGN and thereby dampening the rates of cholesterol and LC-ceramide trafficking from the ER to the Golgi (Capasso et al., 2017). Little is known about the regulation of COPII-dependent bulk membrane trafficking of VLC-ceramides from the ER to the Golgi apparatus.

We, and others, have previously shown that acute depletion of cholesterol from the PM of cultured cells elicits an increase in the synthesis rate of complex VLC-sphingolipids without affecting LC-sphingolipid synthesis (Kim et al., 2023; Perry and Ridgway, 2006). In this study, we showed that that hypothesis is incorrect. We discovered that that VLC-Cer trafficking from the ER to the Golgi is activated by cholesterol depletion and this drives the increase in VLC-SM that is produced by SM synthase activities in the Golgi apparatus.

## Results and Discussion

Acute cholesterol depletion using methyl-beta-cyclodextrin (MβCD) stimulates the synthesis of sphingomyelin (Kim et al., 2023; Perry and Ridgway, 2006). Hydroxypropyl-cyclodextrin (HPCD), which is more selective for cholesterol than MβCD, also caused increased VLC-sphingomyelin synthesis in a dose-dependent manner despite being less effective at cholesterol extraction (Fig. S1). Previously, we showed that only sphingomyelins with very-long-chain (VLC) fatty acids (>20 carbons) increase following cholesterol depletion by MβCD (Kim et al., 2023). We hypothesized that the activity of CerS2, the ceramide synthase responsible for VLC-Cer in the HeLa cell line used in this study (Kim et al 2023) (Fig. S2), is upregulated following cholesterol depletion, thereby increasing VLC-SM production. To test this, we performed a time-course metabolic flux analysis using deuterated sphingosine (d7) and targeted LC-MS/MS to quantify newly synthesized deuterated ceramides and sphingomyelins across various fatty acid chain lengths (C16:0 to C24:1) (Fig. 1A-C).

**Figure 1.**
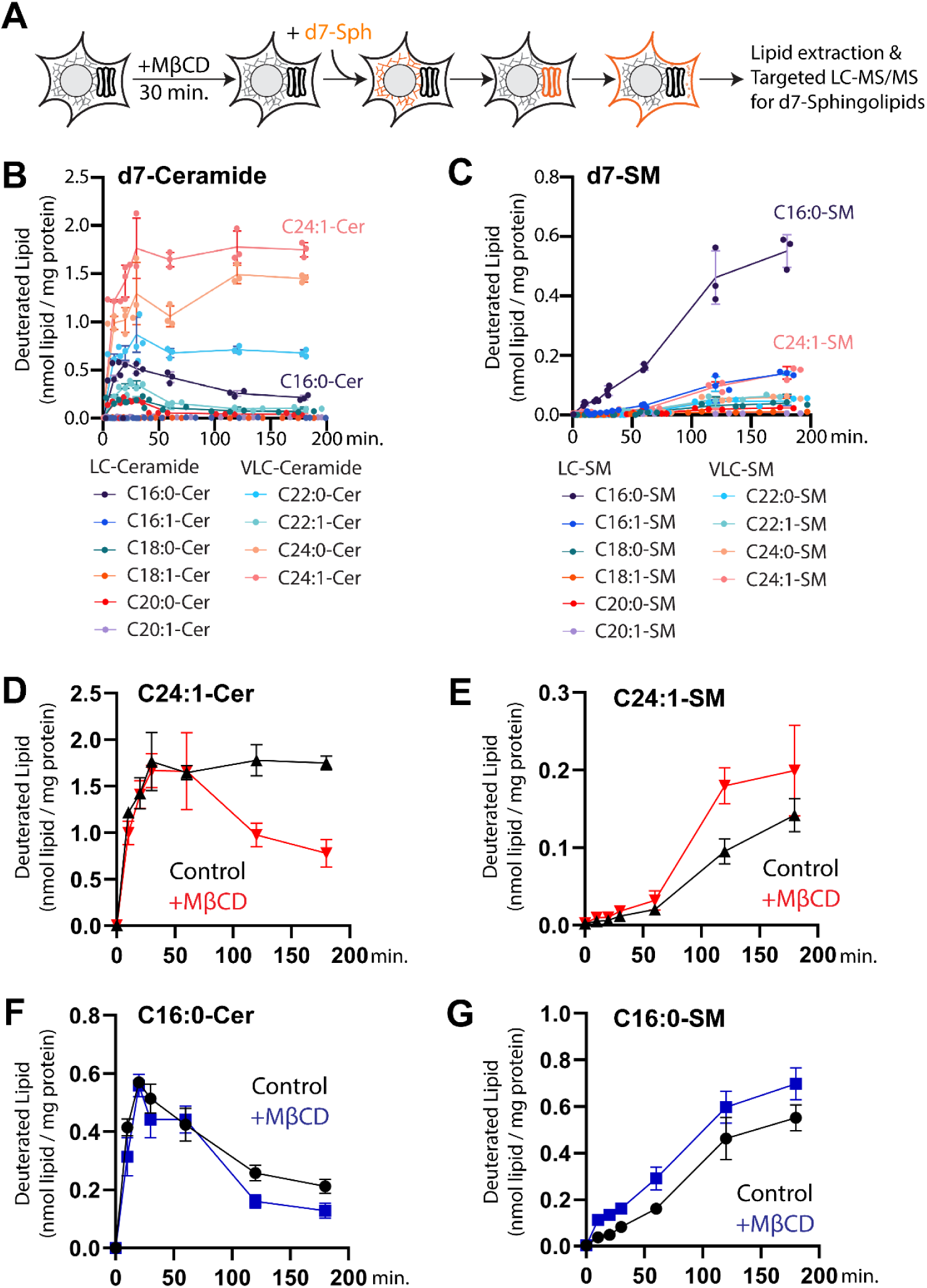
Acute depletion of cholesterol activates sphingomyelin synthesis, but not ceramide synthesis. Sphingolipid metabolism was assessed by metabolic flux analyses using deuterated sphingosine (d7-Sph) and targeted lipid analysis using LC-MS/MS. **A**, Experimental protocol. Cells were treated with DMEM with or without MβCD for 30 minutes, then incubated with the substrate d7-Sph up to 3 hours and lipids were analyzed using LC-MS/MS. **B** and **C**, Time course analyses of ceramide synthesis (B) and sphingomyelin synthesis (C). The chain length of N-linked fatty acids (from 16 to 24 carbons) and the number of double bonds in the fatty acids (0 or 1 double bond) were used to distinguish individual ceramide and SM species. The most abundant VLC species were C24:1-Cer and C24:1 SM and the most abundant LC species were C16:0-Cer and C16:0-SM. We assumed that these 4 species are representative of all newly synthesized sphingolipids for VLC and LC groups. **D and E**, MβCD-treatment increased consumption of C24:1-Ceramide and concomitant increase in C24:1-sphingomylein synthesis. **F** and **G**, C16:0-Cer and C16:0-SM synthesis were not affected by cholesterol depletion.

We compared ceramide synthesis in control and cholesterol-depleted cells. Most abundant LC and VLC sphingolipids (C16:0 and C24:1) were presented. We observed no differences at early time points (<1 hour) after cholesterol extraction (Fig. 1D, F). However, from 2 hours, the amounts of VLC-Cer species began to decrease (Fig. 1D), and there was a concomitant increase in the rate of VLC-SM synthesis (Fig. 1E). The amounts of LC-Cers and LC-SMs changed only modestly (Fig. 1F, G). This suggests that the conversion of VLC-ceramide to VLC-SM was enhanced by cholesterol depletion, even though VLC-Cer synthesis was unchanged. Considering that sphingomyelin synthases (SMS1 and SMS2) reside in the *trans*-Golgi, *trans*-Golgi network (*TGN*), and/or PM (Pothukuchi et al., 2021; Sokoya et al., 2022), and that VLC-SM synthesis relies on COPII-dependent ER-to-Golgi transport (Kim et al., 2023; Loizides-Mangold et al., 2012), these data suggest that cholesterol depletion favors increased VLC-Cer trafficking from the endoplasmic reticulum to the Golgi via a COPII-dependent pathway. Interestingly, VLC-Cer accumulates faster than LC-Cer, but the final product VLC-SM is synthesized slower than LC-SM. This apparent preference to synthesize LC-SM over VLC-SM may occur because CERT-dependent trafficking delivers LC-Cer directly to the *trans*/TGN where SMS1 and SMS2 reside. This CERT-dependent pathway is possibly faster than COPII-dependent transport and intra-Golgi trafficking required for VLC-SM synthesis.

After excluding the possibility that VLC-Cer synthesis was increased by cholesterol depletion (Fig. 1D), we hypothesized that increased SM synthesis after cholesterol depletion is due to enhanced trafficking of VLC-Cer from ER to the Golgi apparatus via the COPII pathway. To correlate sphingolipid trafficking and metabolism analyses, we performed pulse-chase assays using photoactivatable clickable sphingosine (pacSph), which enabled us to visualize labeled lipids *in situ* and to align trafficking information with rates of metabolic conversions that were determined by flux analysis. Cells were incubated with pacSph, then the lipids were extracted. The relative amounts of labeled ceramides, SMs and glucosylceramides were resolved and quantified by thin layer chromatography (Fig. S1A). Cholesterol depletion reduced pacCer and increased pacVLC-SM (Fig. S1B-E), indicating agreement between the deuterated sphingosine and pacSph labeling approaches, allowing us to align trafficking data with metabolic flux values of d7-Sph-containing sphingolipids (Fig. 1). To examine the distribution of pacSph-containing sphingolipids (pacSphingosine) between the ER and Golgi apparatus, cells were incubated with pacSph for 5 to 45 minutes, fixed, and pacSphingolipid was then conjugated *in situ* with fluorescent azide using ‘Click chemistry’ (Fig. 2A). The location of pacSphingolipid adducts in cells was determined using fluorescence microscopy. Accordingly, we measured rates of accumulation of pacSphingolipid within a masked area delimited by the immunofluorescence signal of GM130, a resident of early Golgi cisternae (Fig. S3). Under control conditions, pacSphingolipid adducts were observed in the perinuclear regions (Fig. 2B) and exhibited colocalization with GM130 signals (Pearson’s correlation coefficient, R = 0.5; Fig. 2D). In cholesterol-depleted cells, the intensity of perinuclear pacSphingosine adducts increased, and the colocalization with GM130 signals was enhanced (R = 0.7; Fig. 2D). Importantly, the quantity of pacSphingolipid associated with each cell was similar to control cells, indicating that cholesterol depletion did not enhance the uptake of pacSph (Fig. 2E), but rather enhanced the accumulation of pacSphingolipid-containing adducts in the Golgi.

**Figure 2.**
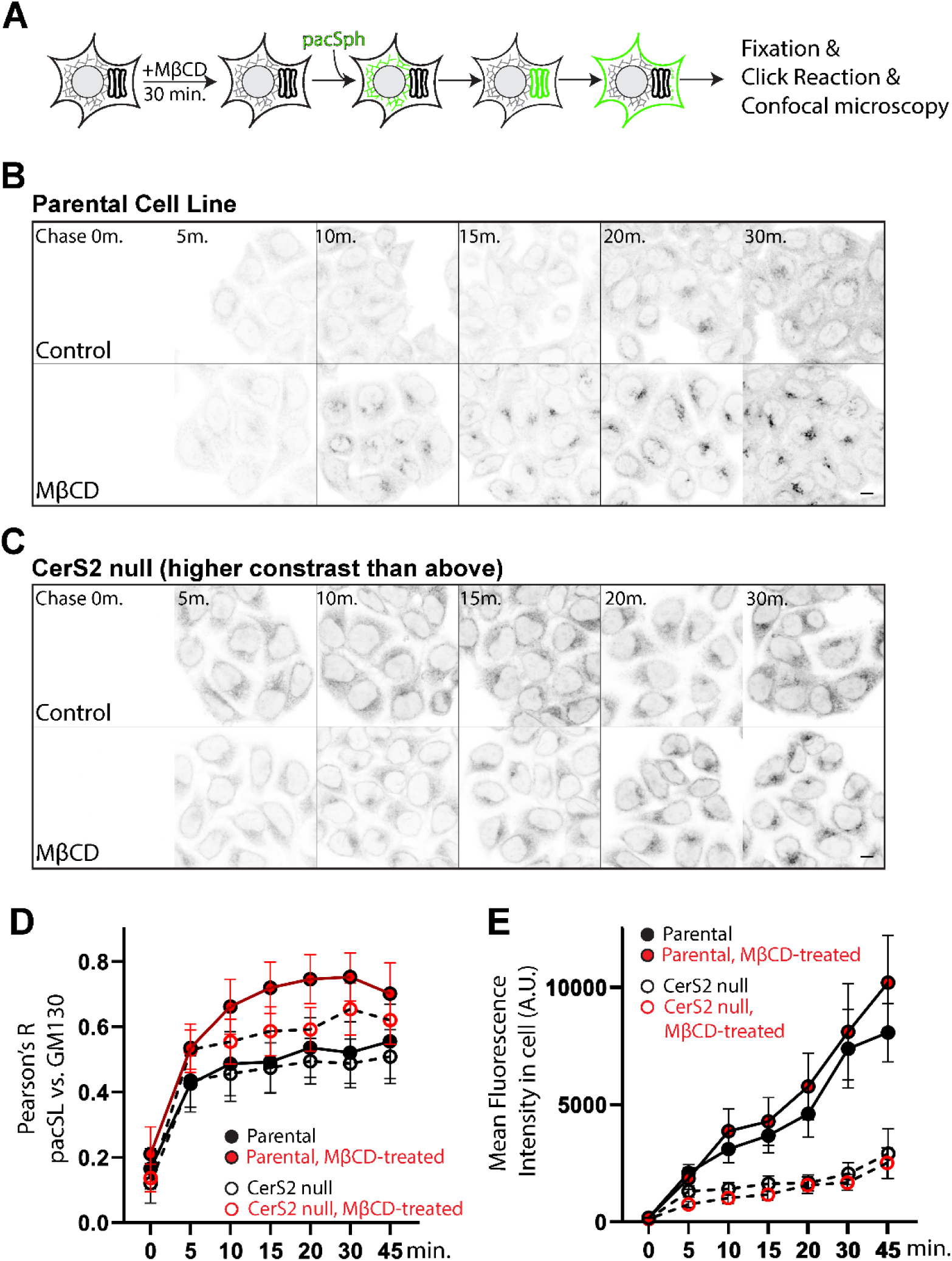
Acute cholesterol depletion increases VLC-Ceramide trafficking from the ER to the Golgi apparatus. Sphingolipid trafficking in cells was assessed by metabolic labeling and *in situ* pacSphingolipid localization. **A**, Experimental protocol. Cells were treated with MβCD for 30 minutes, then incubated with pacSph for the indicated amounts of time prior to fixing cells. pacSphingolipids were conjugated to a fluorophore using click chemistry and visualized by fluorescence microscopy. Anti-GM130 antibody was used to identify early-Golgi compartments by immunofluorescence microscopy. **B and C**, Time course analyses of pacSph adduct localization in WT (parental) (**B**) and CerS2 null (**C**) cells. MβCD-treated cells showed more prominent Golgi-localized pacSphingolipid adducts in CerS2 null cells compared to parental cells. Note that the contrast of the micrographs of CerS2 cells was increased 3-fold (high contrast (3X)) because of the low abundance of pacSphingolipid adducts in these cells. Figure S2 provides an example of the Golgi mask that was determined by anti-GM130 immunofluoresence microscopy. **D,** Colocalization analysis (Pearson’s R) between Golgi marker (GM130) and pacSphingolipid adducts in cells. Pearson’s R values were plotted. More than 500 cell images collected from three independent experiments were used to calculate standard deviation (error bars) and mean values (circles). **E**, The total signal of pacSphingolipid adducts in parental and CerS2 null cells. The data indicate that MβCD-treated cells incorporate pacSph equivalently to control cells and that CerS2 null cells incorporate pacSph at much lower levels than parental cells. The scale bars indicate 10μm.

We addressed if the fluorescence of pacSphingolipid adducts in the Golgi was due to incorporation into pacCer and complex sphingolipids that are made in the Golgi. We utilized CerS2 null cells that produce significantly lower amounts of VLC-sphingolipids (Fig. S2). The mean fluorescence intensities of pacSphingolipid adducts in CerS2 null cells were significantly lower than parental control cells (less than 20% of parental cells) (Fig. 2E), suggesting that pacSph is not retained in cells unless converted to ceramides and that the major lipid signals observed in parental cells primarily originate from ceramides and other complex sphingolipids. It is unlikely that complex sphingolipids, such as SM and hexosyl-ceramide, contribute significantly to the fluorescence concentrated in the Golgi, as experiments using deuterated sphingosine indicated that the primary products identified at the assessed time points (10-120 minutes) were ceramides (Fig. 1B, C). The pacSphingolipid fluorescence signal observed within 5-30 minutes of incubation with pacSph, therefore, is predominantly derived from pacCers.

We investigated whether the cholesterol-dependent changes in pacCer accumulation at the Golgi apparatus were caused by increased amounts of VLC-pacCer rather than LC-pacCer. The results of pacSphingolipid trafficking assays in CerS2 null cells, which produce lower than 5% amounts of VLC-Cer of parental cells (Fig. S2), showed that pacCer accumulation in the perinuclear regions of cells was similar in CerS2 null and parental cells (Fig. 3C). Considering the total fluorescence intensity of pacSphingolipid adducts in the null cells was lower than 20% of the parental cells, a small amount of LC-Cer is trafficked to the Golgi apparatus in CerS2 null cells (Fig. 2D). Cholesterol depletion did not significantly change total cell pacSphingolipid fluorescence in CerS2 null cells (Fig. 2C), and these cells showed a smaller increase in colocalization value of pacSphingolipid adducts (Pearson’s R value, 0.6) with the Golgi marker. Thus, we conclude that VLC-ceramide trafficking from the endoplasmic reticulum to the Golgi apparatus is upregulated when cholesterol is depleted.

**Figure 3.**
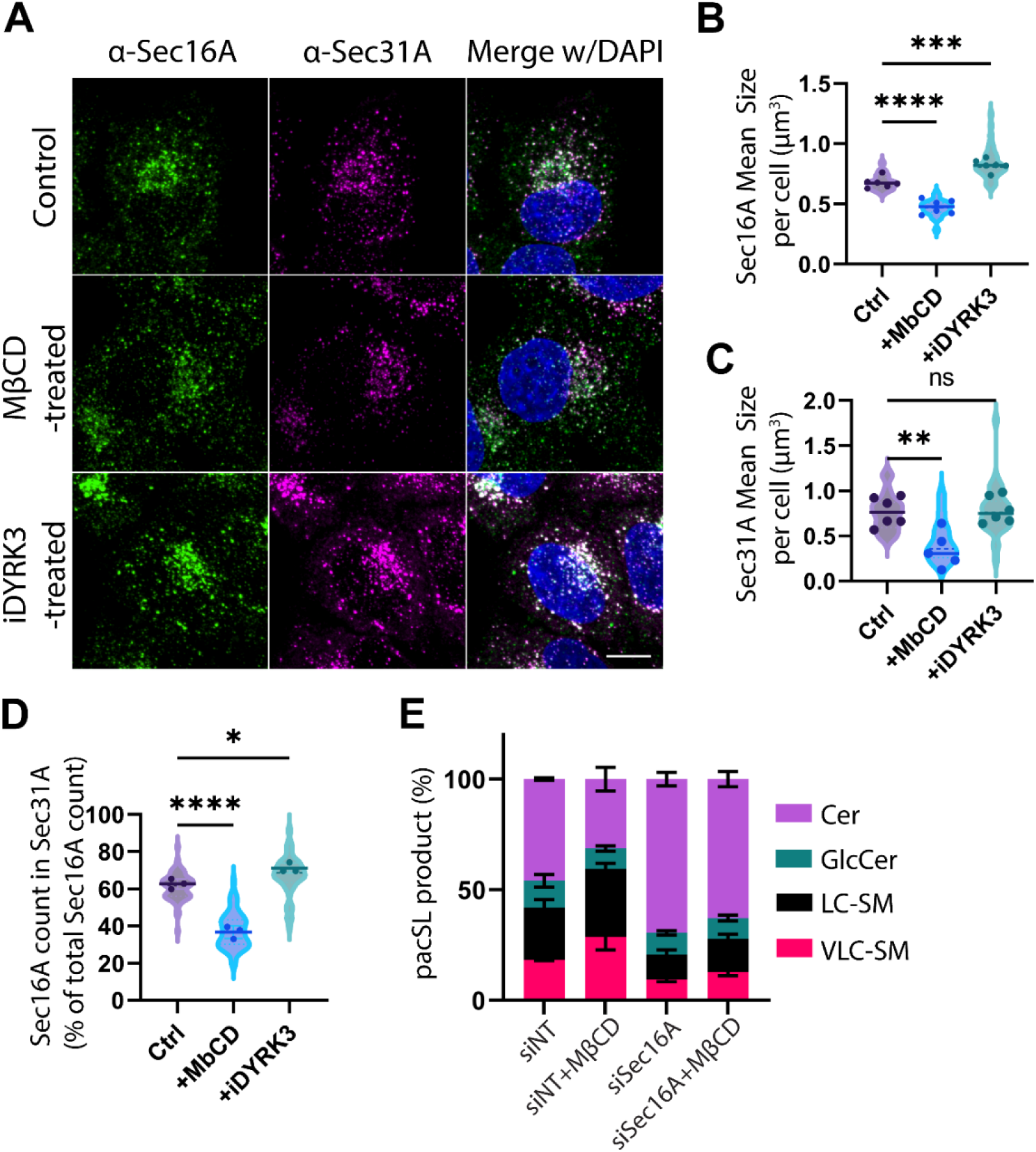
Cholesterol depletion alters the coatomer II ER exit site network. **A**, Immunofluorescence microscopy analysis of Sec16A and Sec31A puncta volumes at the ER exit site. Sec16A (green) and Sec31A (magenta) were visualized by immunofluorescence microscopy. Cells were treated with culture medium (Control), MβCD-containing medium (MβCD-treated), or iDYRK3, then fixed and processed for immunofluorescence. The scale bars indicate 10μm. **B** and **C**, Quantification of mean volume sizes of Sec16A (**B**) and Sec31A (**C**) puncta. Voxels of anti-Sec16A or anti-Sec31A fluorescence in a given cell were used to quantify puncta volumes. Six independent experiments were conducted and mean values (dots) and individual data (violin) are displayed. One-way ANOVA analyses were conducted with mean values from independent experiments. Datasets with less than 0.05 p-value of ANOVA were subjected to Šídák’s multiple comparisons (p-value >0.05, ns; < 0.05, *; < 0.01, **; < 0.001, ***; < 0.0001, ****). **D**, Quantification of the proportions of Sec16A and Sec31A double-positive ERESs in the indicated conditions. Three independent experiments were performed. Data were presented as B and C. **E**, Sphingolipid metabolic analysis of Sec16A-depleted cells. Cells were treated with siRNA for 72 hours, then pacSph pulse-chase assays were conducted. The means and standard deviations of three independent experiments are indicated.

We previously reported that perturbing membrane association of COPII coat proteins by expression of GTP-locked SAR1(H79G) reduced the synthesis of VLC-SM but did not affect synthesis of LC-SM (Kim et al., 2023), leading us to speculate that VLC-Cer is exported from the ER at COPII ER exit sites (ERES). We aimed to see if cholesterol depletion affects ERES by examining the appearance of the COPII-associated proteins, Sec16A and Sec31, by immunofluorescence. Sec16A binds multiple COPII components, though it has not been reported to directly bind to the ER membrane. In cultured *Drosophila melanogaster* cells, Sec16 forms a gel-like structure termed a ‘Sec body’ that forms upon nutrient starvation (Zacharogianni et al., 2014). In HeLa cells growing in nutrient-replete medium, Sec16A exists as a condensate at ERES whose size correlates inversely with rates of protein ER-to-Golgi trafficking (Gallo et al., 2023). Therefore, we speculated that cholesterol depletion elicits a decrease in the size of the Sec16A condensate, and we tested this by anti-Sec16A immunofluorescence analysis of cells subjected to cholesterol depletion (Fig. 3). The size of the Sec16A condensate was previously shown to be enlarged by treatment of cells with an inhibitor of a protein kinase, DYRK3 (GSK626616, iDYRK3, 2.5 µM for 2 hours), and we confirmed that observation (Fig. 3A) (Gallo et al., 2023). We observed that cholesterol depletion led to a reduction in the sizes of both Sec16A and Sec31A puncta (Fig. 3A-C). The proportion of Sec16A and Sec31A double-positive puncta decreased from 62% to 39% after MβCD treatment (Fig. 3D), suggesting that cholesterol is either required for the formation of a complete ERES-COPII network or that cholesterol facilitates release of Sec31A from the ERES, for example, via increased vesicular export. The GSK626616 inhibitor did not elicit any changes in sphingolipid metabolism, indicating that DYRK3-mediated control of ERES dynamics is independent of sphingolipid metabolism and trafficking pathways.

To assess the role of Sec16A in sphingolipid metabolism, we measured pacSph metabolism in cells with SEC16A depletion via siRNAs. Results indicated that Sec16A depletion reduced the synthesis of VLC-SM and increased the amounts of VLC-Cer (Fig. 3E), as we observed previously when COPII activity was disrupted by the expression of a mutant, dominant inhibitory Sar1 GTPase protein (Kim et al., 2023). Furthermore, the upregulation of VLC-SM synthesis elicited by cholesterol depletion was absent in Sec16A-knockdown cells, highlighting Sec16A’s critical role in VLC-Cer metabolism. The efficiency to decrease Sec16A protein levels by siRNA treatment was confirmed (Fig. S5).

To explore the impact of cholesterol depletion on COPII-dependent protein trafficking, we used the Retention-Using-Selective-Hook (RUSH) (Boncompain et al., 2012) system for analyzing fluorescence protein-tagged proteins released from the ER. We focused on glycosylphosphatidylinositol-anchored proteins (GPI-AP), the transferrin receptor 1 (TfR1), and TNF-α (Fig. S3). Our findings showed no significant changes in the rates of ER-to-Golgi trafficking or export from the Golgi of these reporter proteins, suggesting that cholesterol depletion enhances COPII-dependent lipid trafficking, but not COPII-dependent protein trafficking, from the ER to the Golgi apparatus.

Our study suggests that VLC-sphingolipids can be selectively exported from the ER by the COPII network independently of changes in COPII-mediated secretory protein export. This pathway is upregulated upon cholesterol depletion and results in increased SM synthesis in the Golgi apparatus and restoration of plasma membrane homeostasis (Kim et al., 2023). The specificity for VLC-ceramide trafficking and metabolism versus LC-ceramides results simply from the parallel CERT-dependent transport pathway that operates at the ER-Golgi MCS independently of COPII (See Fig. 1 of Kim and Burd, 2023) (Kim and Burd, 2023).

Recent studies show that ERES can be specialized to export different types of protein cargo, and we suggest that sphingolipid export may similarly rely on unique aspects of COPII dynamics to export VLC-ceramide (Phuyal and Farhan, 2021). TANGO1 is a COPII-interacting protein that is crucial for transporting pro-collagen (large cargo) from the ER to the Golgi apparatus by facilitating the concentration of ERES components to the ER and recruiting ERGIC membranes to supply membrane for COPII vesicle formation (Santos et al., 2015; Saxena et al., 2024). Weigel et al (2021) have noted that COPII ERES are pleomorphic networks of tubules emanating from the ER network with a collar of COPII subunits at the base of the tubule and extending to the ERGIC (Weigel et al., 2021). This structure could facilitate sphingolipid transport either by fusing with the ERGIC membrane and/or concentrating ceramide in the ERES membrane network. The increase in SM production in response to cholesterol depletion is caused by an increase in the amount of ceramide that is exported from the ER, rather than changes in ERES number or an increase in the rate of ER-to-Golgi transport of COPII carriers. This observation suggests that ceramide may be concentrated in ER membrane because of cholesterol depletion. Consistent with this, *in vitro* studies of ceramide, phospholipid and cholesterol packing in synthetic vesicles show that ceramide forms an ordered domain in phospholipid-containing vesicles, and it can displace cholesterol from sphingolipid-cholesterol rich domains (Megha and London, 2004). Reduced association of ceramide and cholesterol in the ER membrane may be sufficient to concentrate ceramide and facilitate its rapid transport to the ERGIC. Further research will be necessary to test if sphingolipid-specific or protein trafficking-independent membrane trafficking exists in cells.

## Methods

### Cell lines and chemicals

HeLa cells from CSL cell line service (300194/p772_HeLa) were grown in DMEM (11965-092, Gibco) supplemented with 10% fetal bovine serum (FBS) (16140-071, Gibco) and maintained at 37°C with 5% CO_2_, unless otherwise noted. All cell lines were grown in humidified incubator at 37°C with 5% CO_2_ and were routinely screened for mycoplasma contamination using a mycoplasma detection kit (30-1012K, ATCC).

SGPL1 null HeLa was described in previous research from our lab (Deng et al., 2018). CerS2 null HeLa and CerS2-SGPL1 double null HeLa were generated by following a previously published CRISPR/Cas9 gene-targeting protocol (Ran et al., 2013). Two guide RNA (targeting sequences: 5’-tcggatgtaattggcaaacc and 5’-ctcttccgattacctgctgg) were each inserted into the CRISPR/Cas9-expressing plasmid, pX459 2.0 mammalian expression vector (Addgene plasmid # 62988). Methyl-β-cyclodextrin (MβCD, 377110050) and (2-hydroxypropyl)-β -cyclodextrin (HPCD, H31133.06) were purchased from Millipore Sigma. GSK-626616 was purchased from MedChemExpress (iDYRK3, HY-105309).

### Transfection, plasmids and siRNA

Lipofectamine 2000 (11668-019, Invitrogen) was used for transient transfection of cells with plasmids in accordance with the manufacturer’s instructions (For RUSH assay, 100 ng GalT-BFP and 200 ng pCMV-Str-KDEL_SS-SBP-GFP-GPI plasmid DNA with 2.5 µl of Lipofectamine 2000 in 1 ml). mApple-TNFα-RUSH and mApple-TfR1-RUSH were purchased from Addgene and used for RUSH imaging (Weigel et al., 2021). All analyses of transfected cells were done within 24 hours after transfection. siRNA sequence targeting Sec16A was 5’-CCAGGTGTTTAAGTTCATCTA (Piao et al., 2017). Lipofectamine RNAiMAX (13778075, Invitrogen) was used for 72 hours, following the manufacturer’s protocol.

### pacSph pulse-chase experiments

Photoactivatable-clickable-sphingosine (pacSph) was purchased from Avanti Polar Lipids (900600). Solvents were purchased as following; Hexane (HX0295-1, Sigma), ethyl acetate (9282-33, JT Baker), chloroform (9180-01, JT Baker), methanol (9070-05, JT Baker), acetate (695092, Sigma). TLC was purchase from Sigma (EM1.05721.0001). Metabolic labeling experiments performed as described previously (Kim et al., 2023). Briefly, cells were grown in 10 cm dishes and treated accordingly. 3 µM pacSph was given to cells in the medium (DMEM) for 1 hour in the incubator at 37°C with 5% CO_2_. Cells were washed with DMEM and incubated with DMEM 1 hour in the incubator at 37°C with 5% CO_2_. Cells were harvested at 4°C. Cells were washed with PBS and stored at −80°C until lipid extraction. Lipids were extracted (Haberkant et al., 2016) and reacted with Click reagents as described (Gerl et al., 2016). Extracted pacSphingolipids were reacted with 3-azido-7-hydroxycoumarine (CLK-FA04701, Jena Bioscience). Coumarine-conjugated pacSphingolipids were applied to TLC plates and separated by solvent A (chloroform:methanol:water:acedic acid in volume ratio 65:25:4:1), then by solvent B (ethylacedate:hexane in 1:1 volume). Separated fluorescent lipids were visualized and quantified by BioRad Molecular Imager ChemiDoc XRS+ system and ImageLab Software (BioRad).

### Cholesterol measurement

Cholesterol levels were measured by using a cholesterol assay (A12216, Invitrogen) and the manufacturer’s protocol. Cholesterol esterase was not added in the assay to measure the level of cholesterol, not cholesterol ester.

### RUSH live cell imaging and image analysis

For RUSH live cell imaging, 250 000 cells were seeded and transfected in glass bottom dishes (P35G-1.5-14-C, MatTek). To remove traces of biotin from the growth media, cells were incubated after transfection with RUSH cargo plasmids in the presence of avidin (2 μg/ml). In the following day, cells were incubated in the absence or presence of 5 mM MβCD in plain DMEM for 120 minutes at 37 °C with 5% CO_2_. Cells were imaged in DMEM at 37 °C with 5% CO_2_, and the RUSH cargo was released by the presence of biotin (0.1 mg/ml). The fluorescence micrographs were captured using a SoRa CSU-W1 (Yokogama) spinning disk confocal workstation (Nikon) based on an inverted microscope (Nikon Ti2-E) and ORCA-FusionBT back-thinned camera (Hamamatsu). Images were captured using 60× oil lens (N.A. 1.4), immersion oil for 22°C or 37°C and NIS-Elements Advanced Research Package (Nikon). All micrographs under analysis were a single confocal slice.

The images were analyzed using FIJI and in-house macros (Schindelin et al., 2012). To analyze the trafficking of RUSH cargo toward Golgi apparatus, co-transfected cells were selected to quantify the fluorescence intensity in the Golgi marker (GalT-BFP). The mean intensity of Golgi-localized RUSH fluorescence was normalized from 0 (minimum mean intensity) to 1 (maximum mean intensity of the Golgi marker throughout time points). Time-dependent changes of single-cell data were plotted by Normalized fluorescence intensity (F.I.) in GalT-BFP over time.

### Immunofluorescence and image analysis

Antibodies were purchased from BD Biosciences (Sec31A, BDB612351; GM130, 610823), and from Invitrogen (Sec16A, PA5-52182). DAPI was purchased from Invitrogen (34580). Goat anti-Rabbit secondary antibody (488) (A-11008) and goat anti-mouse antibody (A-11029) were purchased. Cells were seeded and grown overnight in DMEM with 10% FBS at 37°C with 5% CO_2_. For MβCD treatment in Fig. 4, cells were treated with 5 mM MβCD for 30 minutes, washed with DMEM, and incubated with DMEM for 90 minutes. For iDYRK3 treatment, cells were treated with 5 µM iDYRK3 in DMEM for 120 minutes. At the end of treatment, cells were fixed with 4% paraformaldehyde in PBS, diluted from 16% paraformaldehyde (Electron Microscopy Sciences, 15710), for 15 minutes at room temperature. Cells were permeabilized by 0.1% saponin in PBS for 5 minutes at room temperature. Cells were blocked by 5% FBS in PBS for 1 hour. Primary antibodies (a-Sec16A and a-Sec31A) were diluted in 1:200 in 5% FBS and used overnight at 4°C. Secondary antibodies and Hoechst were used for 1 hour at room temperature. The fluorescence micrographs were captured using a SoRa CSU-W1 with an inverted microscope (Nikon Ti2-E) and ORCA-FusionBT back-thinned camera (Hamamatsu). Z-stacks with 0.2 µm increment, covering 9 µm from the bottom of the cell were obtained. Three emission channels were used; one to receive Alexa 488 signals conjugated with the secondary antibody targeting anti-Sec16A antibody; one for Alexa 546 targeting Sec31A; one for DAPI signals.

For the analysis of Sec16A and Sec31 immunofluorescence signals using ImageJ and FIJI (Schindelin et al., 2012), immunofluorescence signal was defined as top 2.5% intensity of whole confocal image fluorescence. The single threshold value among every confocal image under analysis was calculated and used for the subsequent analysis. Plugins called “3D OC Options” and “3D Objects Counter” were used to count the puncta of Sec16A and Sec31A per cell and calculate the volume of them. To calculate Sec16A signals associated with Sec31A, double-channel-positive pixels were selected and used to calculate the puncta volume; puncta bigger than 10 voxels (above minimum confocal resolution 200 nm x 200 nm x 200 nm) were defined as true double-positive puncta. The number of double-positive puncta was used to calculate Sec16A count in Sec31A (% of total Sec16A count).

### In situ lipid visualization by Click reaction and image analysis

For MβCD treatment in Fig. 3, cells were treated with 5 mM MβCD for 30 minutes, washed with DMEM, and incubated with pacSph (1 µM final) in DMEM for designated timepoints in the Fig 3. For in situ click reaction, Click-iT Cell Reaction Buffer Kit (C10269) was purchased from Life Technologies. Alexa Fluor 647 azide was purchased from Invitrogen (A10277). After staining for primary (GM130), secondary antibodies, and DAPI, cells were treated with Click reagents, according to the manufacturer’s protocol from Life Technologies (C10269). Cells were washed and fluorescence was visualized by confocal fluorescence microscopy (SoRa CSU-W1, Nikon Ti2-E, ORCA-FusionBT back-thinned camera), using three channels (633, 546, and 488 channels).

Image analysis was performed by using ImageJ and FIJI plugins (Schindelin et al., 2012). For the analysis of pacSph pulse-chase images, three emission channels were analyzed; one to receive Alexa633 signals conjugated with pacSphingolipids; one to receive Alexa 488 signals conjugated with the secondary antibody targeting anti-GM130 antibody; one to receive DAPI signals for nuclei staining. DAPI signal was used to count the cell number and cell area. “Find Maxima” process was used to locate local maxima DAPI signal and to identify single cells region of interest (ROI). Mean fluorescence intensity of cell in the 633 channel was measured by finding mean values in each cell ROI. The plugin “Coloc 2” was used to calculate Pearson’s R values between pacSphingolipid and GM130 signals for each cell ROI.

### Sphingolipid metabolic analysis using liquid chromatography-mass spectrometry - mass spectrometry

For lipidomic and sphingolipid metabolic flux analysis, internal standards including SPLASH® Lipidomix® Mass Spec Standard (330707), SphingoSplash^TM^ (330734), and C15 Ceramide-d7 (d18:1-d7/15:0) (860681P) were purchased from Avanti Polar Lipids. To identify fragmentation patterns and retention times of targeted lipids, CER Internal Standard Mixture - Ultimate SPLASH (330823), Deuterated Ceramide LIPIDOMIX® Mass Spec Standard (330713X), Ceramide LIPIDOMIX® Mass Spec Standard (330712X) were purchased from Avanti Polar Lipids. HPLC- or LCMS-grade organic solvents including methanol (C959B47), acetonitrile (AX0156), 2-propanol (1027814000), methyl-t-butyl ether (MtBE, MX0826) and chloroform (C990U23) were purchased from Millipore Sigma. Ammonium acetate, 7.5 M solution was purchased from MP Biomedicals (0219875980).

Cells were seeded in 10 cm tissue culture dishes. At approximately 90% confluency, cells were harvested by lifting and washing with PBS. Cell pellets were flash-frozen in liquid nitrogen at stored at −80°C until lipid extraction. On the day of lipid extraction, cells were resuspended in 400 µL of ice-cold PBS and homogenized by tip sonicator in the cold room (4°C). 1 µL of lysates were used for quantifying protein levels by using Bradford protein assay. 400 uL of Lysates were centrifuged at 100,000 x g for 1 hour at 4°C using Beckman Coulter Optima MAX-XP Ultracentrifuge. The supernatant was discarded, and the membrane pellets were resuspended in 1 mL of MtBE:MeOH (3:1 vol/vol) solution that contains internal standards (Splash Lipidomix I, and SphingoSplash I). Glassware was used for all lipid handling. The membrane resuspension was vortexed for 45 minutes at 4°C, then sonicated in the ice-filled water bath for 15 minutes. The membrane solution was centrifuged at 1000 x g for 5 minutes at 4°C. The supernatant was transferred to a new glass tube. 650 µL of H_2_O:MeOH (3:1, vol/vol) containing 0.2% formic acid was added to the tube and the mixed by vortexing for 1 minute. Tubes were centrifuged at 1000 x g at 4°C for 15 minutes to separate the phase. The upper phase that contains most of MtBE were collected into a new glass tube. To the leftover lower phase, 650 µL of MtBE (0.2% formic acid) was added and the tube was mixed. Tubes were centrifuged at 1000 x g at 4°C for 15 minutes to separate the phase. The upper phase was transferred to the first MtBE phase above. Solvents were dried by streaming nitrogen gas. Lipids were resuspended in 50 µL of Chloroform:MeOH (1:4, vol/vol) and transferred to Agilent glass vials and inserts. Glass vials were stored at −20°C for no more than 2 days until used for mass spectrometry.

Agilent 6546 LC/Q-TOF and InfinityLab Poroshell 120 EC-C18 (695575-902) column were used to analyze lipidome and deuterated lipids. 10 microliters of samples were loaded using Agilent 1290 Infinity II LC. The mass detection range was set between 300 to 1700 m/z for positive and negative ions. Column pumps were set at 60°C. Auto MS/MS mode was used. Acquisition rate was 4 spectra / second for MS and MS/MS. For precursor selection, 3 max precursor per cycle was used. Active exclusion was used, excluding precursors after 2 spectra, releasing after 0.2 minutes. Q-TOF Gas temp was set to 325°C, and sheath gas temp was set to 275°C. Fragmentor voltage was set to 275°C. Mobile phase A (60% acetonitrile, 40% H_2_O, 7.5 mM ammonium acetate) and phase B (90% isopropanol, 10% acetonitrile, 7.5 mM ammonium acetate) were used to separate lipids. Gradients started at 85% A and decreased to 70% A over 2 minutes, then to 52% A over 30 seconds. The gradient slowly decreased to 18% A over 12.5 minutes, then 1% in 1 minute. The gradient was held for 4 minutes. The gradient was restored to 85% A for 5 minutes for washing. The next sample was run from the first gradient. Lipids were identified using MS-Dial (Tsugawa et al., 2020). Lipid identities were manually curated by using fragmentation patterns and retention times. The amounts were normalized to the amount of internal standards.

### Immunoblot analysis

Cells were treated (siRNA) as described above. Whole-cell lysates were prepared in a sample buffer containing 250 mM Tris-HCl, pH 6.8, 50% glycerol, 10% sodium dodecyl sulfate, 10% β-mercaptoethanol, 0.025% bromophenol. Protein contents were measured by Coomassie-protein quantification solution (23200, Pierce). Proteins were resolved by SDS-PAGE and transferred to nitrocellulose blots. The blots were blocked by 5% FBS in TBS with 0.1% tween 20 (TBST). Anti-Sec16A antibody was purchased from Invitrogen (Sec16A, PA5-52182) and used in 1:2000 dilution in TBST. After overnight, blots were washed with TBST and incubated with Horse-Radish Peroxidase (HRP)-conjugated secondary antibody (1:10,000, 7074S, Cell Signaling Technology) for 1 hour. Blots were washed and incubated with HRP substrates (170-5060, Bio-Rad). Chemiluminescence from blots and Ponceau-stained blots were imaged by Molecular Imager ChemiDoc XRS+ System (Bio-Rad).

### Statistical analyses and data presentation

GraphPad Prism 9 was used for statistical analyses and presentation of quantitative data. Student’s t-test was used for two-sample comparisons. One-way ANOVA was used to compare more than two conditions. When p-value of ANOVA was lower than 0.05, post-hoc analysis (Dunnett’s multiple comparison test) was performed (Fig. 3). P-values were indicated as following: * p<0.05, ** p<0.01, *** p<0.001, **** p<0.0001.

## Supplemental material

**Summary**: Figures S1-S3. The experimental results from the investigation on the effects of cholesterol depletion on sphingomyelin synthesis, the sphingolipidome, pacSphingolipid trafficking, and ER-to-Golgi lipid and protein trafficking.

**Figure S1.**
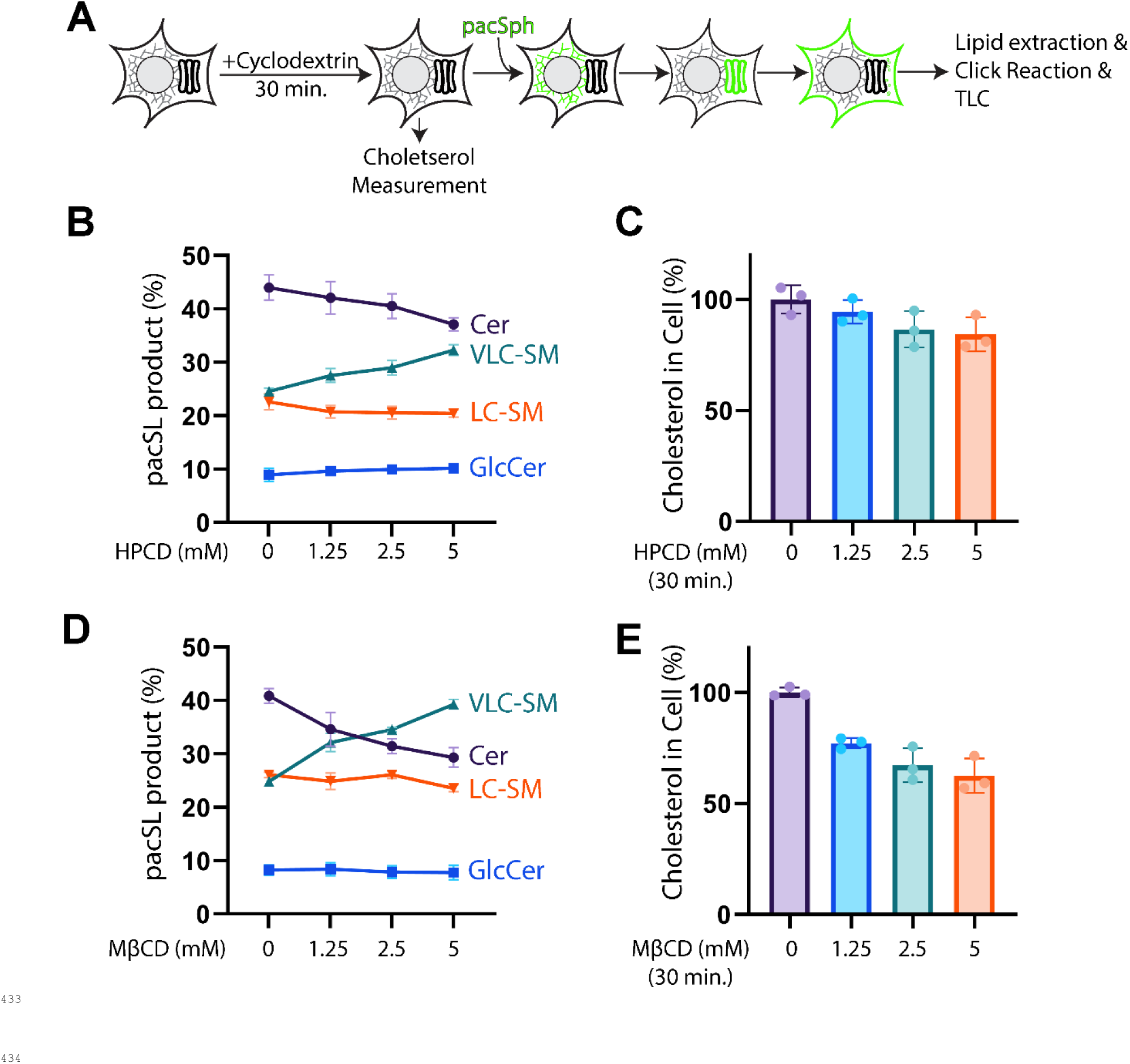
Very-Long-Chain sphingomyelin synthesis is increased upon cholesterol depletion. **A**, Experimental protocol. For metabolic labeling, cells were first incubated with cyclodextrin (MβCD or HPCD) for 30 min. The MβCD was then removed and pacSph labeling (60 min) was initiated. After the pulse period, the media were replaced with fresh media to begin the chase period (60 min). For assessing cholesterol levels, cells were incubated with cyclodextrin (MβCD or HPCD) for 30 min and were used for cholesterol measurement. pacSph pulse-chase analysis of sphingolipids in cells treated with varying amounts of HPCD (**B**) or MβCD (**D**). The relative amounts of four pacSph-containing products, Cer, glucosylceramide (GlcCer), VLC-SM and LC-SM were quantified and mean values and standard deviations of three independent experiments are shown. **C** and **E** show the cholesterol content of cells incubated with the indicated amounts of HPCD or MβCD, respectively, for 30 minutes. The individual measurement (dots), mean values (bar size) and standard deviations (error bars) of three independent experiments are shown.

**Figure S2.**
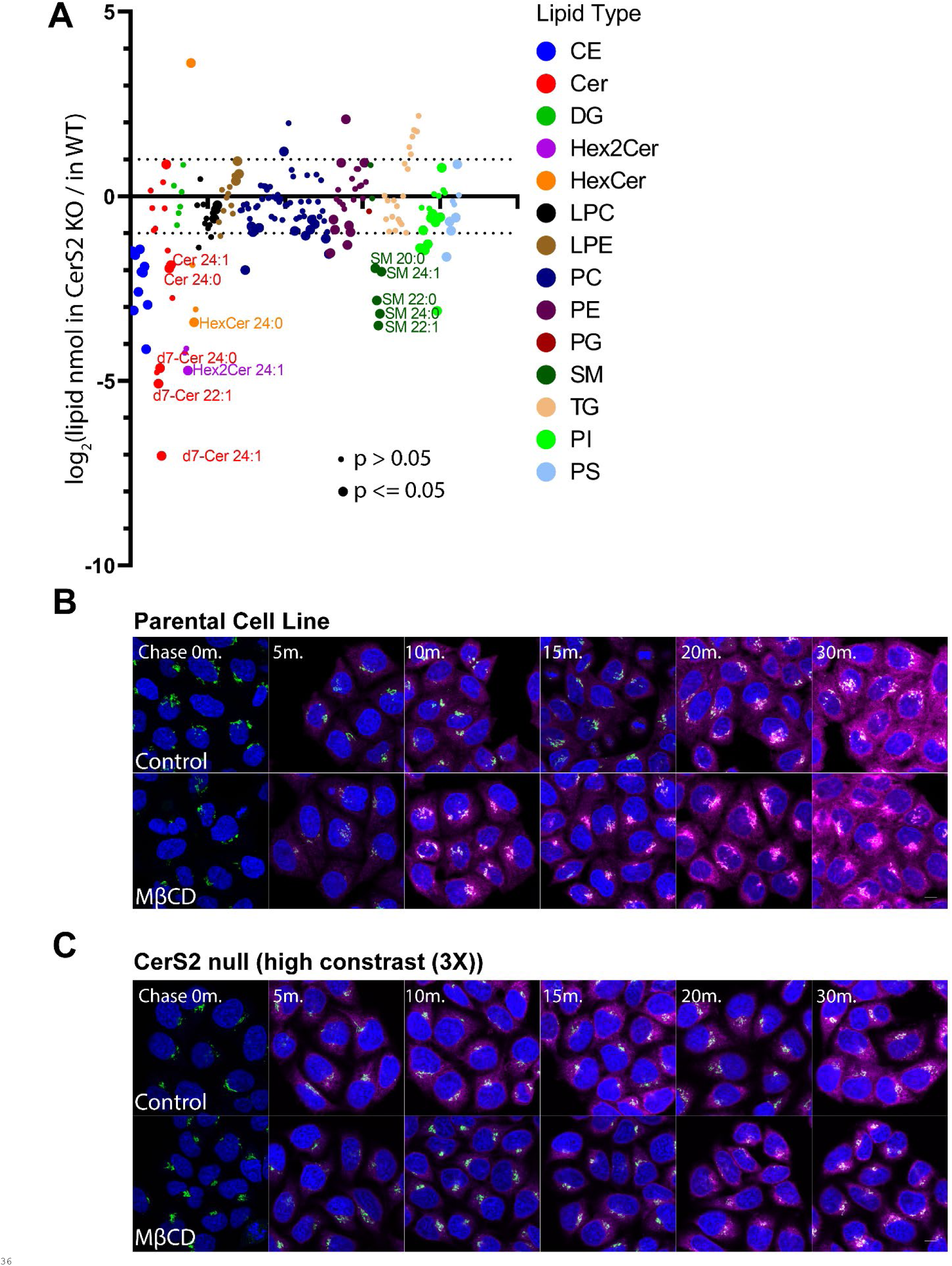
Quantitative lipidomics and pacSphingolipid trafficking analyses. **A**, The lipidome analysis of CerS2 null cells. WT and CerS2 KO cells were incubated with deuterated sphingosine (d7) for 30 minutes, and total lipidome and deuterated lipids were analyzed by quantitative LC/MS/MS. Cells were grown, treated as above, and harvested three times independently. Lipids were quantified by internal standards with known concentration (nmol unit) and normalized by mg of proteins in the sample. Lipid types were color-coded (CE, cholesteryl ester; Cer, ceramide; DG, diacylglycerol; Hex2Cer, 2-hexosylceramide; HexCer, hexosylceramide; LPC, lysophosphatidylcholine; LPE, lysophosphatidylethanolamine; PC, phosphatidylcholine; PE, phosphatidylethanolamine; PG, phosphatidylglycerol; SM, sphingomyelin; TG, triacylglycerol; PI, phosphatidylinositol; PS, phosphatidylserine). The mol ratio of quantified lipids were log-2-scaled. P-values from Student’s t-ttest were calculated using lipid quantities (nmol) and displayed as bigger dots (p<=0.05) or smaller dots (p>0.05). **B-C,** Fluorescence micrographs of *in situ* pacSphingolipid adducts (magenta) and anti-GM130 immunostaining (green). The scale bars indicate 10μm. The micrographs show anti-GM130 immunostaining that was used to mask the Golgi in the micrographs shown in Figure 2B and 2C. To visualize weaker signals of pacSphingolipid adducts in CerS2 null cells, the threshold for maximum intensity was lowered for pacSphingolipid adduct signals (high contrast, (3X)).

**Figure S3.**
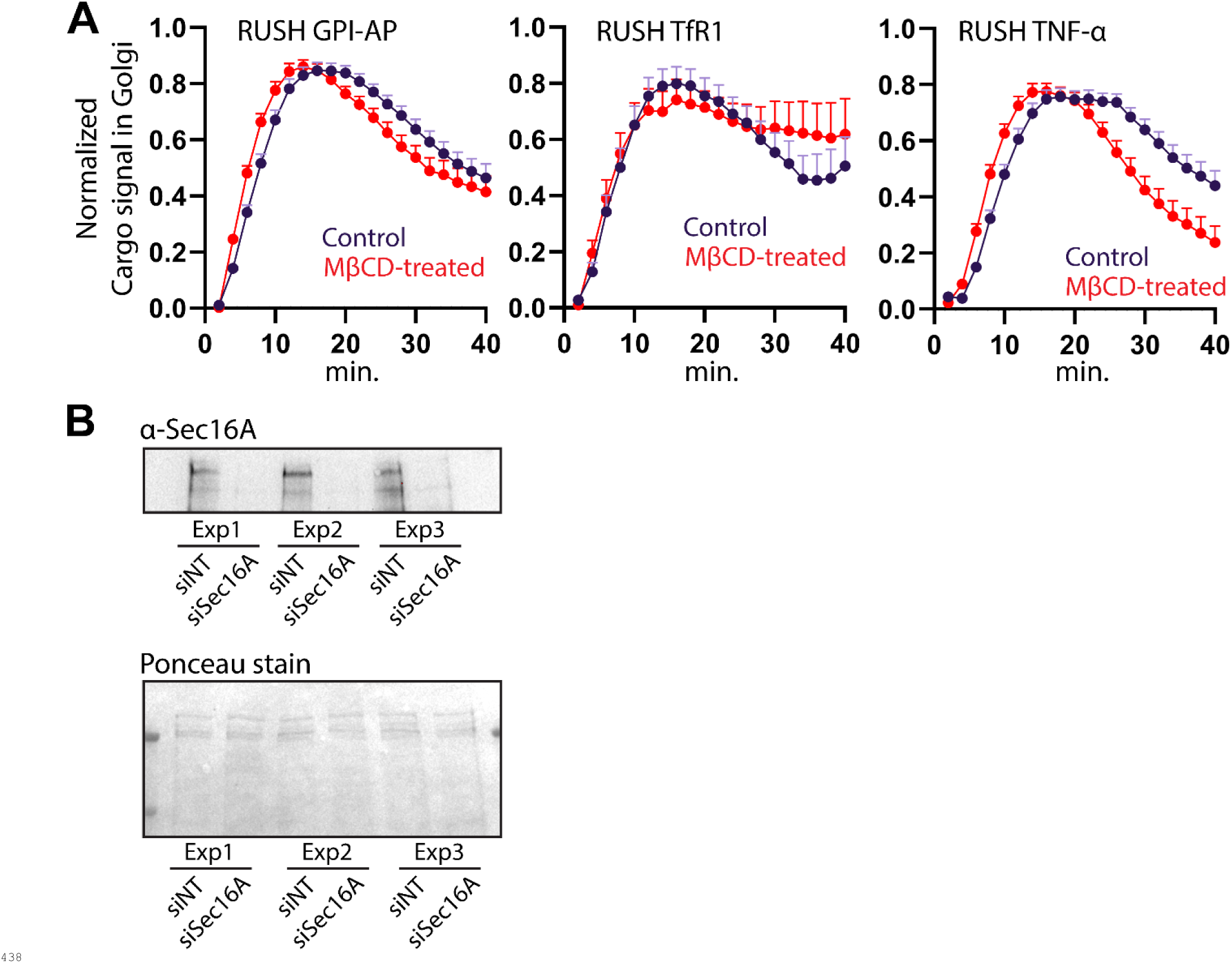
Analysis of ER-to-Golgi protein trafficking and Sec16A protein levels. **A**, ER-to-Golgi protein trafficking was assessed by RUSH cargo trafficking assay of three independent reporter proteins. Live cell imaging was used to measure the rates of appearance of cargo trafficking reporters after release from the ER by addition of biotin. The normalized fluorescence intensity from RUSH cargo proteins in a mask of a co-expressed Golgi resident protein (GalT-BFP) was determined. Time-dependent changes of single-cell data were plotted by Normalized Cargo signal in Golgi (GalT-BFP mask). Error bars indicate the 95% confidence interval collected from single cell data, derived from 50∼200 cells. Cells expressed RUSH GPI-AP (GFP-tagged glycosylphosphatidylinositol anchored protein), RUSH mApple-tagged transferrin receptor 1 (RUSH TfrR1) and RUSH mApple-tagged tumor necrosis factor-α (RUSH TNF-α). **B**, Confirmation of Sec16A depletion by siRNAs. Immunoblotting of endogenously expressed Sec16A in three independent experiments. Ponceau staining was used to gauge the amounts of protein in each lane.

## Data availability statement

All data are available in the published article and its supplemental material.

## Acknowledgements

We are grateful to colleagues for critical reading and suggestions. We thank Julia von Blume, Thomas Melia and Xiaolei Su (Department of Cell Biology, Yale School of Medicine, New Haven, CT, USA) for sharing reagents and facilities. This work is supported by funds from the National Institute of General Medical Sciences of the National Institutes of Health under award numbers GM144096 and GM095766-08S1 (CGB).

## Author Contributions

CGB and YK conceptualized and CGB supervised this study. CGB secured research funding. YK performed all experiments and prepared the initial drafts of the text and figures. JP performed protein trafficking experiments using RUSH constructs. YK and CGB reviewed and edited the manuscript.

## Abbreviations

Cer: ceramide
CerS: ceramide synthase
CERT: ceramide transfer protein
COPII: coatomer II
d7-SM: 7-deuterated sphingomyelin
d7-Sph: 7-deuterated sphingosine
DYRK3dual: specificity tyrosine-phosphorylation-regulated kinase 3
ER: endoplasmic reticulum
ERES: endoplasmic reticulum exit site
ERGIC: endoplasmic reticulum-Golgi intermediate compartment
GPI-AP: glycosylphosphatidylinositol-anchored protein
GSL: glycosphingolipid
HPCD: hydroxypropyl-cyclodextrin
LC: long-chain
LTP: lipid transfer protein MCS membrane contact site
MβCD: methyl-beta-cyclodextrin
OSBP: oxysterol binding protein
pac: photoactivatable clickable
pacCer: photoactivatable clickable ceramide
pacSph: photoactivatable clickable sphingosine
pacSphingolipid: photoactivatable clickable sphingolipid
pacVLC-SM: photoactivatable clickable very-long-chain sphingomyelin
PI4P: phosphatidyl-inositol 4-phosphate
SM: sphingomyelin
SPTLC: serine-palmitoyl-transferase
TfR1: transferrin receptor 1
TGN: trans-Golgi network
TNF-*α*: tumor necrosis factor alpha
VLC: very-long-chain

## References

Balla, A., G. Tuymetova, A. Tsiomenko, P. Varnai, and T. Balla. 2005. A plasma membrane pool of phosphatidylinositol 4-phosphate is generated by phosphatidylinositol 4-kinase type-III alpha: studies with the PH domains of the oxysterol binding protein and FAPP1. Mol Biol Cell. 16:1282–1295.

Boncompain, G., S. Divoux, N. Gareil, H. de Forges, A. Lescure, L. Latreche, V. Mercanti, F. Jollivet, G. Raposo, and F. Perez. 2012. Synchronization of secretory protein traffic in populations of cells. Nat Methods. 9:493–498.

Breslow, D.K., S.R. Collins, B. Bodenmiller, R. Aebersold, K. Simons, A. Shevchenko, C.S. Ejsing, and J.S. Weissman. 2010. Orm family proteins mediate sphingolipid homeostasis. Nature. 463:1048–1053.

Capasso, S., L. Sticco, R. Rizzo, M. Pirozzi, D. Russo, N.A. Dathan, F. Campelo, J. van Galen, M. Holtta-Vuori, G. Turacchio, A. Hausser, V. Malhotra, I. Riezman, H. Riezman, E. Ikonen, C. Luberto, S. Parashuraman, A. Luini, and G. D’Angelo. 2017. Sphingolipid metabolic flow controls phosphoinositide turnover at the trans-Golgi network. EMBO J. 36:1736–1754.

Cingolani, F., A.H. Futerman, and J. Casas. 2016. Ceramide synthases in biomedical research. Chem Phys Lipids. 197:25–32.

Deng, Y., M. Pakdel, B. Blank, E.L. Sundberg, C.G. Burd, and J. von Blume. 2018. Activity of the SPCA1 Calcium Pump Couples Sphingomyelin Synthesis to Sorting of Secretory Proteins in the Trans-Golgi Network. Dev Cell. 47:464–478 e468.

Gallo, R., A.K. Rai, A.B.R. McIntyre, K. Meyer, and L. Pelkmans. 2023. DYRK3 enables secretory trafficking by maintaining the liquid-like state of ER exit sites. Dev Cell. 58:1880–1897 e1811.

Gerl, M.J., V. Bittl, S. Kirchner, T. Sachsenheimer, H.L. Brunner, C. Luchtenborg, C. Ozbalci, H. Wiedemann, S. Wegehingel, W. Nickel, P. Haberkant, C. Schultz, M. Kruger, and B. Brugger. 2016. Sphingosine-1-Phosphate Lyase Deficient Cells as a Tool to Study Protein Lipid Interactions. PLoS One. 11:e0153009.

Haberkant, P., F. Stein, D. Hoglinger, M.J. Gerl, B. Brugger, P.P. Van Veldhoven, J. Krijgsveld, A.C. Gavin, and C. Schultz. 2016. Bifunctional Sphingosine for Cell-Based Analysis of Protein-Sphingolipid Interactions. ACS Chem Biol. 11:222–230.

Hanada, K., K. Kumagai, S. Yasuda, Y. Miura, M. Kawano, M. Fukasawa, and M. Nishijima. 2003. Molecular machinery for non-vesicular trafficking of ceramide. Nature. 426:803–809.

Kim, Y., and C.G. Burd. 2023. Lipid Sorting and Organelle Identity. Cold Spring Harb Perspect Biol. 15.

Kim, Y., G. Mavodza, C.E. Senkal, and C.G. Burd. 2023. Cholesterol-dependent homeostatic regulation of very long chain sphingolipid synthesis. J Cell Biol. 222.

Lange, Y., M.H. Swaisgood, B.V. Ramos, and T.L. Steck. 1989. Plasma membranes contain half the phospholipid and 90% of the cholesterol and sphingomyelin in cultured human fibroblasts. J Biol Chem. 264:3786–3793.

Levine, T.P., and S. Munro. 2002. Targeting of Golgi-specific pleckstrin homology domains involves both PtdIns 4-kinase-dependent and -independent components. Curr Biol. 12:695–704.

Loizides-Mangold, U., F.P. David, V.J. Nesatyy, T. Kinoshita, and H. Riezman. 2012. Glycosylphosphatidylinositol anchors regulate glycosphingolipid levels. J Lipid Res. 53:1522–1534.

Megha, and E. London. 2004. Ceramide selectively displaces cholesterol from ordered lipid domains (rafts): implications for lipid raft structure and function. J Biol Chem. 279:9997–10004.

Perry, R.J., and N.D. Ridgway. 2006. Oxysterol-binding protein and vesicle-associated membrane protein-associated protein are required for sterol-dependent activation of the ceramide transport protein. Mol Biol Cell. 17:2604–2616.

Phuyal, S., and H. Farhan. 2021. Want to leave the er? We offer vesicles, tubules, and tunnels. J Cell Biol. 220:2–4, pmid = 33999114.

Piao, H., J. Kim, S.H. Noh, H.S. Kweon, J.Y. Kim, and M.G. Lee. 2017. Sec16A is critical for both conventional and unconventional secretion of CFTR. Sci Rep. 7:39887.

Pothukuchi, P., I. Agliarulo, M. Pirozzi, R. Rizzo, D. Russo, G. Turacchio, J. Nuchel, J.S. Yang, C. Gehin, L. Capolupo, M.J. Hernandez-Corbacho, A. Biswas, G. Vanacore, N. Dathan, T. Nitta, P. Henklein, M. Thattai, J.I. Inokuchi, V.W. Hsu, M. Plomann, L.M. Obeid, Y.A. Hannun, A. Luini, G. D’Angelo, and S. Parashuraman. 2021. GRASP55 regulates intra-Golgi localization of glycosylation enzymes to control glycosphingolipid biosynthesis. EMBO J. 40:e107766.

Ran, F.A., P.D. Hsu, J. Wright, V. Agarwala, D.A. Scott, and F. Zhang. 2013. Genome engineering using the CRISPR-Cas9 system. Nat Protoc. 8:2281–2308.

Santos, A.J., I. Raote, M. Scarpa, N. Brouwers, and V. Malhotra. 2015. TANGO1 recruits ERGIC membranes to the endoplasmic reticulum for procollagen export. Elife. 4.

Saxena, S., O. Foresti, A. Liu, S. Androulaki, M. Pena Rodriguez, I. Raote, M. Aridor, B. Cui, and V. Malhotra. 2024. Endoplasmic reticulum exit sites are segregated for secretion based on cargo size. Dev Cell. 59:2593–2608 e2596.

Schindelin, J., I. Arganda-Carreras, E. Frise, V. Kaynig, M. Longair, T. Pietzsch, S. Preibisch, C. Rueden, S. Saalfeld, B. Schmid, J.Y. Tinevez, D.J. White, V. Hartenstein, K. Eliceiri, P. Tomancak, and A. Cardona. 2012. Fiji: an open-source platform for biological-image analysis. Nat Methods. 9:676–682.

Sokoya, T., J. Parolek, M.M. Foged, D.I. Danylchuk, M. Bozan, B. Sarkar, A. Hilderink, M. Philippi, L.D. Botto, P.A. Terhal, O. Makitie, J. Piehler, Y. Kim, C.G. Burd, A.S. Klymchenko, K. Maeda, and J.C.M. Holthuis. 2022. Pathogenic variants of sphingomyelin synthase SMS2 disrupt lipid landscapes in the secretory pathway. Elife. 11.

Tsugawa, H., K. Ikeda, M. Takahashi, A. Satoh, Y. Mori, H. Uchino, N. Okahashi, Y. Yamada, I. Tada, P. Bonini, Y. Higashi, Y. Okazaki, Z. Zhou, Z.J. Zhu, J. Koelmel, T. Cajka, O. Fiehn, K. Saito, M. Arita, and M. Arita. 2020. A lipidome atlas in MS-DIAL 4. Nat Biotechnol. 38:1159–1163.

Wang, Y.J., J. Wang, H.Q. Sun, M. Martinez, Y.X. Sun, E. Macia, T. Kirchhausen, J.P. Albanesi, M.G. Roth, and H.L. Yin. 2003. Phosphatidylinositol 4 phosphate regulates targeting of clathrin adaptor AP-1 complexes to the Golgi. Cell. 114:299–310.

Weigel, A.V., C.L. Chang, G. Shtengel, C.S. Xu, D.P. Hoffman, M. Freeman, N. Iyer, J. Aaron, S. Khuon, J. Bogovic, W. Qiu, H.F. Hess, and J. Lippincott-Schwartz. 2021. ER-to-Golgi protein delivery through an interwoven, tubular network extending from ER. Cell. 184:2412–2429 e2416.

Zacharogianni, M., A. Aguilera-Gomez, T. Veenendaal, J. Smout, and C. Rabouille. 2014. A stress assembly that confers cell viability by preserving ERES components during amino-acid starvation. Elife. 3.

